# CD20^+^ natural killer cells are polyfunctional, memory-like cells that are enriched in inflammatory disorders

**DOI:** 10.1101/2025.01.09.632131

**Authors:** Özgür Albayrak, Ergün Tiryaki, Nazan Akkaya, Ali Burak Kızılırmak, Tansu Doran, Gökçe Gökmenoğlu, Muhammed Yüksel, Bürge Ulukan, Mina Üzülmez, Işıl Baytekin, Kemal Soylu, Güneş Esendağlı, Ingrid Meinl, Mesrure Köseoğlu, Burcu Yüksel, Suat Erus, Çiğdem Arıkan, Seçil Vural, Müjdat Zeybel, Aysun Soysal, Edgar Meinl, Atay Vural

## Abstract

While CD20 was initially characterized as a B cell-specific marker, its expression on memory T cells has expanded our understanding of this molecule’s distribution and function. Here, we identify a previously unrecognized CD20-expressing NK cell population and demonstrate its functional significance. CD56+CD20+ NK cells exhibit hallmarks of cellular activation, including elevated NKp46, CD69, and CD137 expression, enhanced proliferative capacity, and increased production of inflammatory cytokines (IFN-γ, GM-CSF, TNF-α, IL-10). Functional analyses revealed enhanced cytotoxicity against K562 targets, correlating with increased expression of cytolytic mediators including granzymes A, B, and K, perforin, FASL, and TRAIL. Single-cell transcriptional profiling demonstrated that MS4A1-expressing NK cells possess a distinct molecular signature characterized by elevated granzyme K expression and memory-like features. These cells preferentially localize to secondary lymphoid organs and accumulate in inflammatory tissues. Notably, CD56+CD20+ NK cells are enriched in multiple inflammatory conditions, including multiple sclerosis, autoimmune hepatitis, hepatitis B infection, hepatocellular carcinoma, and lung cancer. Treatment with rituximab depletes this population, suggesting potential therapeutic implications. Our findings establish CD20+ NK cells as a functionally distinct lymphocyte subset with enhanced effector capabilities and tissue-homing properties, providing new insights into immune regulation in inflammatory diseases.

**One Sentence Summary:** Our study reveals expression of CD20 by NK cells, in relation with enhanced functionality, memory-like features, and inflammation.

## INTRODUCTION

CD20 is expressed abundantly on B cells and depletion of B cells by CD20 monoclonal antibodies proved to be an effective treatment modality in multiple sclerosis (MS), underscoring their pivotal role in MS pathogenesis (*1*). Intriguingly, other B cell-specific treatments like atacicept and belimumab that do not target CD20 have not demonstrated effectiveness in MS, raising the question whether CD20 contributes to MS pathogenesis through mechanisms beyond its presence on B cells (*2*).

In addition to B cells, CD20 is expressed at low levels by a fraction of T cells (*3, 4*). These CD3^+^CD20^+^ lymphocytes were shown to have an active phenotype with strong cytokine secretion upon stimulation, are enriched in type 1 cytotoxic CD8^+^ T cells and have a predominantly memory T cell phenotype (*4, 5*). Moreover, CD20^+^ T cells are also depleted by CD20 monoclonal antibody therapies, and their depletion was shown to ameliorate experimental autoimmune encephalomyelitis independent of B cells (*6–8*). These and other studies demonstrated that CD3^+^CD20^+^ cells are clinically relevant in several autoimmune disorders like MS, rheumatoid arthritis, Sjögren’s syndrome, psoriasis, and have a role in cancer immunosurveillance (*9, 10*). It remains, however, unknown whether CD20 expression is restricted solely to cells of the adaptive immune response.

In this study, we report natural killer (NK) cells also express CD20 on their surface and thus demonstrate that CD20 expression is not limited to adaptive lymphocytes. In addition to having a more active phenotype like CD3^+^CD20^+^ cells, CD20^+^ NK cells exhibit greater degranulation capacity and cytotoxicity against cancer cells. Importantly, CD56^+^CD20^+^ cells have a higher proliferative capacity and possess memory-like molecular features. We also show that CD56^+^CD20^+^ cells are enriched in tissues, especially in secondary lymphoid organs. Finally, by examining blood and tissue samples from patients with a wide range of disorders, and publicly available single-cell RNA sequencing (scRNAseq) datasets, we show that CD56^+^CD20^+^ cells are involved in the pathogenesis of several autoimmune disorders, HBV infection and cancer.

## RESULTS

### CD20 expression on NK cells

In a cohort of healthy controls (HCs) (mean age 30.3±1.3, F:M=9:5), we observed that 1-4.5% of peripheral blood NK cells express CD20 at a frequency similar to that of CD3^+^CD20^+^ T cells (median, IQR: 3.3, 2.4-3.9 vs 4.4, 3.7-5, respectively) (Figure 1, A and B). CD20 was present on both CD56^bright^ and CD56^dim^ cells, the ratio being higher in the former (median, IQR: 4.2, 3.5-5.1 vs 2.9, 1.9-3.5, respectively) (Figure 1B).

**Figure 1.**
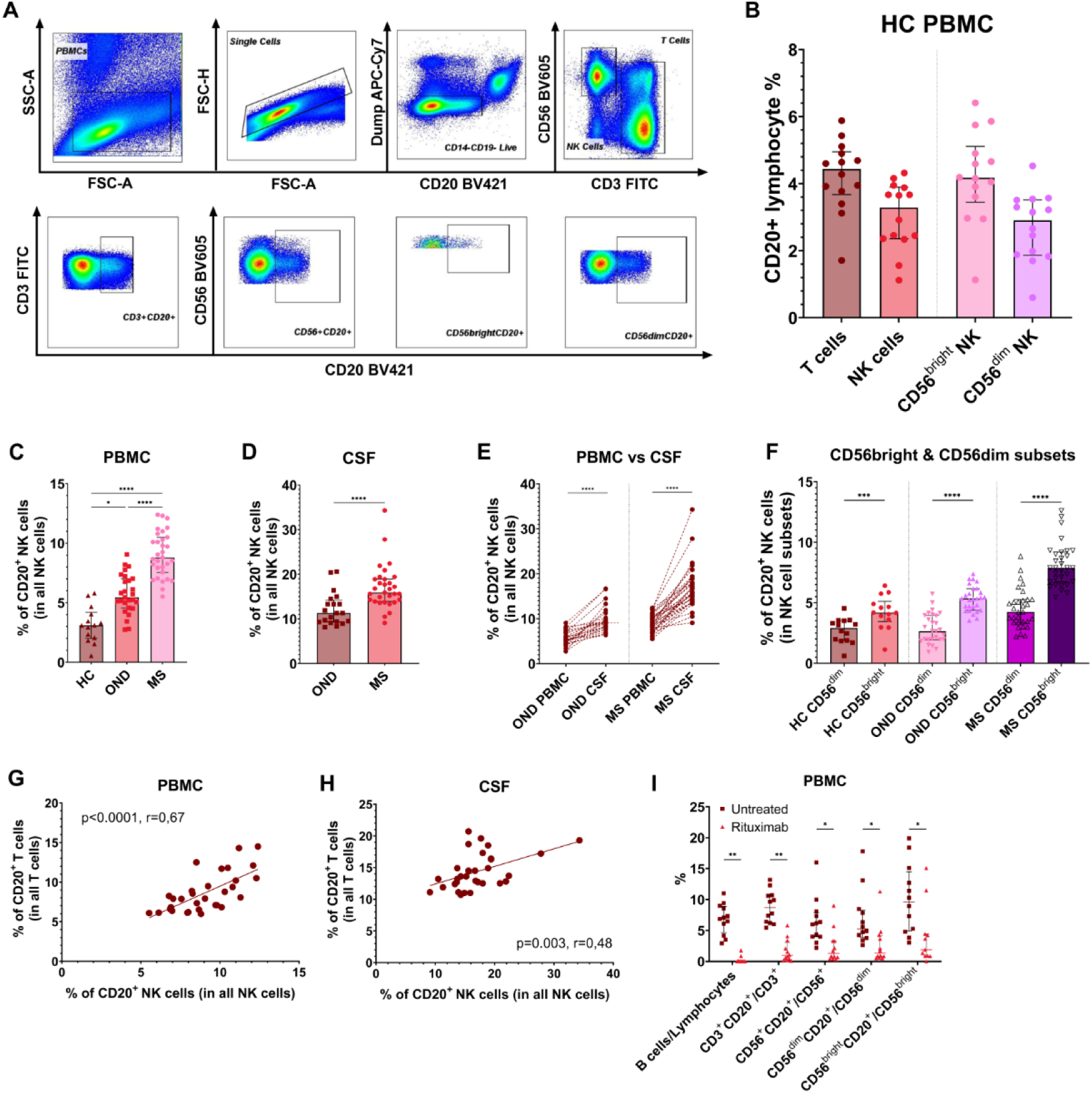
**(A)** Gating strategy of live CD3^−^CD14^−^CD19^−^CD56^+^CD20^+^ cells. **(B)** The frequency distribution of CD20^+^ NK cells and T cells in the blood of healthy volunteers. **(C)** The ratio of CD20^+^ NK cells to all NK cells (CD56^+^CD20^+^ %) is greater in the blood of people with newly diagnosed multiple sclerosis (pwMS, n=31) compared to the patients with other neurological disorders (OND, n=27), and healthy controls (HCs, n=14). P values were calculated using Kruskal-Wallis test with Dunn’s multiple comparisons post hoc correction. **(D)** The ratio of CD56^+^CD20^+^ cells is higher in the CSF of pwMS compared to the patients with other neurological disorders. P value was calculated using two-tailed Mann-Whitney test. **(E)** CSF is enriched for CD56^+^CD20^+^ cells compared to the blood in both patient groups. **(F)** CD56^bright^ NK cells are enriched for CD20^+^ cells in both HCs and neurological patients. P values were calculated using two-tailed Wilcoxon matched-pairs signed rank test. **(G and H)** The frequency of CD56^+^CD20^+^ cells in the peripheral blood (G) and CSF (H) of pwMS are correlated with that of CD3^+^CD20^+^ cells in both tissues. P and r values were calculated by Pearson’s correlation test. **(I)** CD56^+^CD20^+^ cells are depleted alongside with B cells and CD3^+^CD20^+^ cells after rituximab treatment compared to untreated pwMS. *P < 0.05, **P < 0.01, ***P < 0.001, ****P < 0.0001.

### The percentage of CD20^+^ NK cells is higher in the blood and cerebrospinal fluid of people with MS compared to controls

In a previous study the percentage of CD20^+^ T cells was reported to be increased in the peripheral blood and cerebrospinal fluid (CSF) of people with MS (pwMS) and this increase was found to be related to disease severity (*5*). We asked whether the frequency of CD56^+^CD20^+^ cells is also increased in pwMS and answered this question by analyzing the paired peripheral blood and CSF of a cohort of patients with new onset of neurological symptoms, including a demyelinating attack (table S1).

We found that the ratio of CD20^+^ NK cells to all NK cells is significantly higher in pwMS (mean ± SD: 9 ± 1.9) compared to other neurological disorders (OND) (5.7 ± 1.6, p<0.0001) and HCs (3.2 ± 1.5, p<0.0001) in the peripheral blood. Similarly, the percentage of CD20^+^ NK cells was higher in the CSF of pwMS (17 ± 4.9) compared to OND (12.2 ± 3.7, p<0.0001) (Figure 1, C and D). Clinical features of the study participants are shown in Table S1. CD20^+^ NK cells were enriched in CSF compared to peripheral blood in all patients (Figure 1E). CD56^+^CD20^+^ cells were more frequent in the CD56^bright^ population compared to the CD56^dim^ (Figure 1F). In addition, the frequency of CD56^+^CD20^+^ cells was significantly correlated with that of CD3^+^CD20^+^ cells and B cells in pwMS in both the peripheral blood (p<0.0001 and p<0.05, respectively) and CSF (p<0.01 and p<0.01, respectively) (Figure 1, G and H; Figure S1, A and B).

### CD20^+^ NK cells in blood are depleted after rituximab therapy

Both B cells and CD20^+^ T cells are known to be depleted after rituximab infusion and CD3^+^CD20^+^ cells were shown to replenish before B cells (*4*). We compared the ratio of CD56^+^CD20^+^ cells obtained from 12 rituximab treated (median, IQR: 1.3, 0.5-3.3) and 12 untreated (6, 4-7.9) pwMS and found significantly lower CD56^+^CD20^+^ cell frequency in the rituximab group (p<0.05). Moreover, CD56^+^CD20^+^ cell ratio was higher compared to B cells in treated patients, indicating that CD20^+^ NK cells may also repopulate faster than B cells (Figure 1I).

### CD20^+^ NK cells have a more active phenotype and secrete more cytokines

After recognizing the significance of CD20^+^ NK cells in MS, we aimed to elucidate the phenotype and functional characteristics of these cells in both resting conditions and after stimulation.

We found that a higher percentage of CD20^+^ NK cells express the activating receptor NKp46 (p<0.01), and activation markers, CD137 (p<0.01) and CD69 (p<0.001) compared to CD56^+^CD20^−^ cells, whereas the ratio of CD57^+^ (p<0.01) and CD16^+^ (p<0.001) cells was higher among CD20^−^ NK cells (Figure 2A and B). The percentage of NKG2A, NKG2C and KIR expressing cells were similar in both groups (Figure 2C).

**Figure 2.**
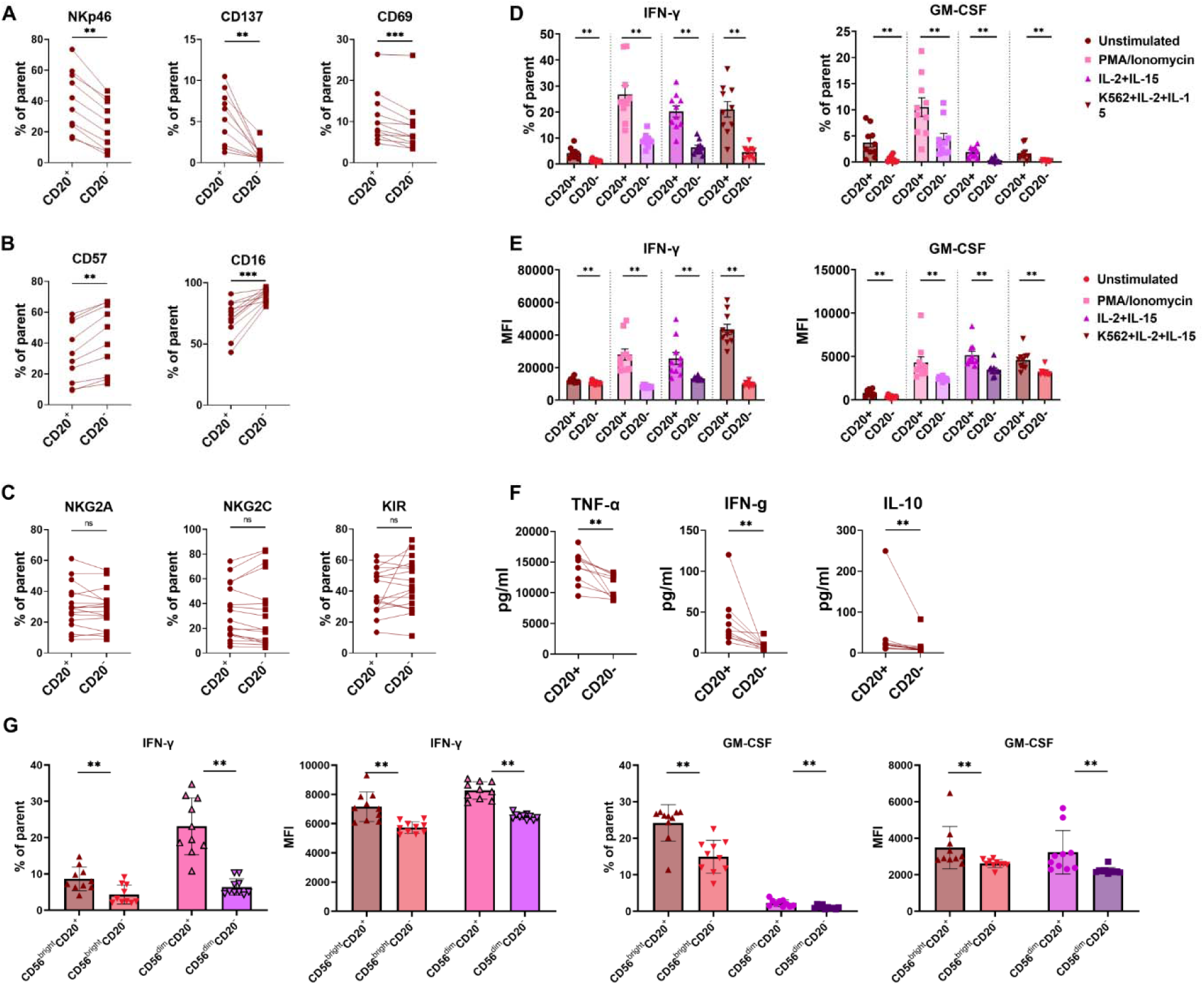
CD20^+^ NK cells have a more active phenotype and secrete more cytokines. **(A and B)** The ratio of NKp46, CD69 and CD137 expressing cells are higher among CD20^+^ NK cells (A), whereas the ratio of CD57^+^ and CD16^+^ cells are higher among CD20^−^ NK cells (B). **(C)** The percentage of NKG2A, NKG2C and KIR expressing cells are similar in both groups. **(D and E)** Intracellular staining and flow cytometry analysis of sorted NK cells from healthy donors revealed that the percentage of cells producing IFN-γ and GM-CSF (D) and the amount produced, as shown by mean fluorescence intensity (MFI) measurements, (E) are higher in CD56^+^CD20^+^ cells after stimulation with PMA/Ionomycin, IL-2 plus IL-15, and K562 cells plus IL-2 plus IL-15. **(F)** Stimulated CD56^+^CD20^+^ cells secreted a higher amount of TNF-α, IFN-γ and IL-10 into the cell culture medium compared to the CD20^−^ NK cells. **(G)** Subgroup analysis of cells stimulated with IL-2 plus IL-15 showed that the ratio and MFI of IFN-γ and GM-CSF expression are higher in both CD56^bright^CD20^+^ and CD56^dim^CD20^+^ cells, compared to their CD20^−^ counterparts. P values were calculated using two-tailed Wilcoxon matched-pairs signed rank test. *P < 0.05, **P < 0.01, ***P < 0.001, ****P < 0.0001.

Next, we assessed sorted NK cells from healthy donors by intracellular staining for their cytokine production capacity under three different stimulation conditions: PMA/Ionomycin, IL-2 plus IL-15, and K562 cells plus IL-2 plus IL-15. The percentage of cells producing IFN-γ and GM-CSF was higher in CD56^+^CD20^+^ cells under all conditions (p<0.01 for all conditions tested for both cytokines) (Figure 2D). The MFI of these cytokines were also higher in CD56^+^CD20^+^ cells (p<0.01 for all conditions tested for both cytokines) (Figure 2E). Moreover, CD56^+^CD20^+^ cells stimulated with K562 cells plus IL-2 and IL-15 secreted more TNF-α (p<0.01), IFN-γ (p<0.01) and IL-10 (p<0.01) into the cell culture medium compared to the CD20^−^ NK cells (Figure 2F).

Subgroup analysis of the CD56^bright^ and CD56^dim^ populations revealed that both the percentage of NK cells expressing IFN-γ and GM-CSF, as well as their MFI levels are higher in CD56^bright^CD20^+^ and CD56^dim^CD20^+^ cells compared to their CD20^−^ counterparts (Figure 2G).

The percentage of CD56^+^CD20^+^ cells among CD56^+^ NK cells and the MFI of CD20 were not changed after 6h under either of the stimulation conditions (Figure S2, A and B).

### CD20^+^ NK cells produce and secrete greater quantities of cytotoxic molecules

Natural killer cells are critical for the elimination of virally infected and cancerous cells. Therefore, we next investigated the production of cytotoxic molecules by CD56^+^CD20^+^ cells during incubation with K562 cancer cells under IL-2 plus IL-15 stimulation. We found that the MFIs of granzyme A (GrA), granzyme B (GrB), perforin and granzyme (GrK) but not granulysin, were higher in CD20^+^ NK cells compared to CD20^−^ NK cells in the resting condition and after stimulation. (Figure 3, A to E).

**Figure 3.**
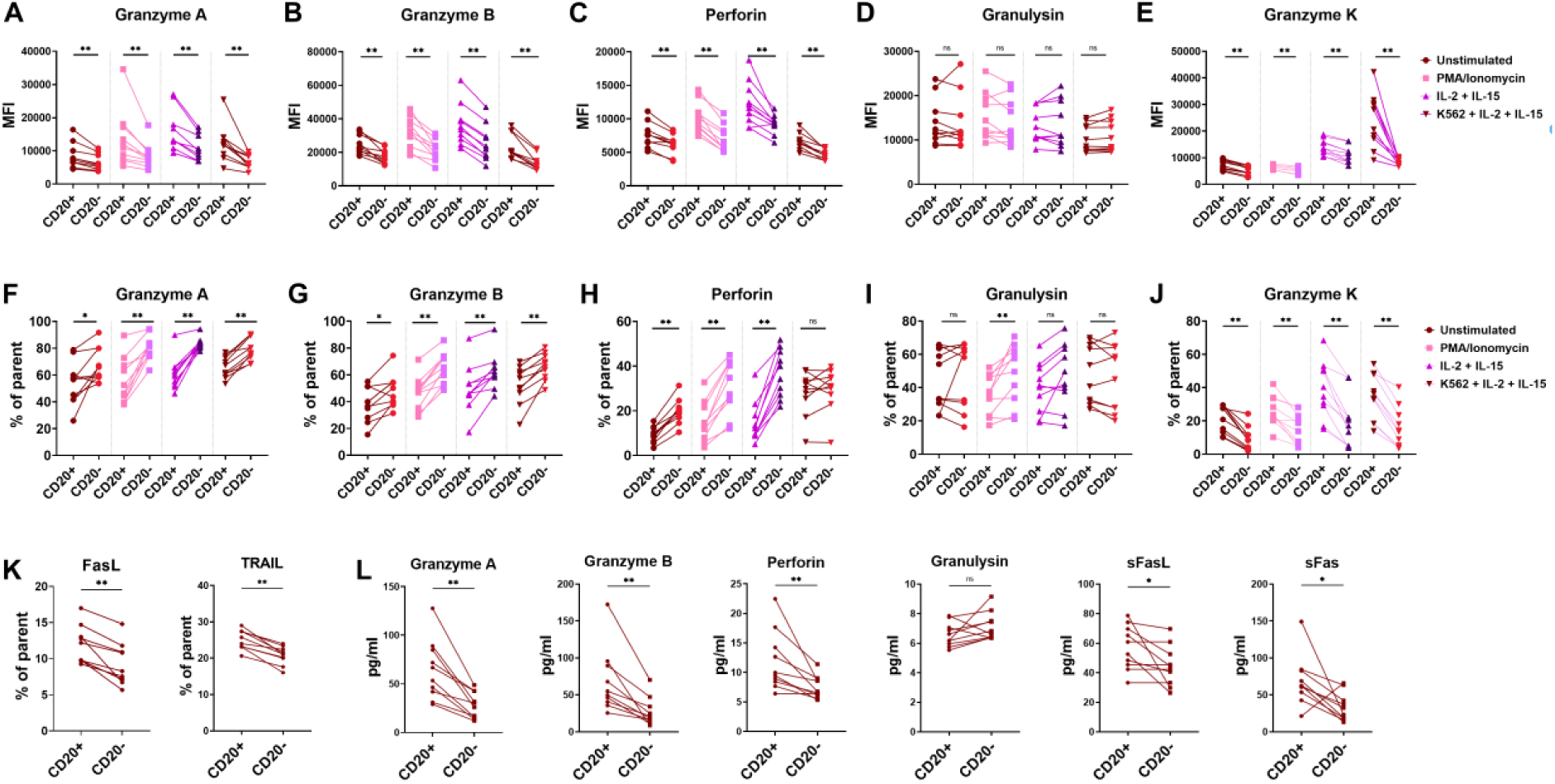
CD20^+^ NK cells produce and secrete cytotoxic molecules more abundantly. **(A to E)** The amount of GrA, GrB, perforin and GrK but not granulysin produced by CD20^+^ NK cells are higher compared to CD20^−^ NK cells, especially after stimulation. **(F to J)** However, only the percentage of GrK expression is higher among CD20^+^ NK cells, whereas the ratio of GrA, GrB, and perforin expressing cells are higher among CD20^−^ NK cells. **(K)** A greater proportion of the CD56^+^CD20^+^ cells express FasL, and TRAIL on the cell surface. **(L)** CD56^+^CD20^+^ cells secrete higher amounts of GrA, GrB, perforin, soluble FasL and soluble Fas into the cell culture medium after stimulation. P values were calculated using two-tailed Wilcoxon matched-pairs signed rank test. *P < 0.05, **P < 0.01.

Conversely, the percentage of cells expressing GrA, GrB, and perforin was lower in CD56^+^CD20^+^ cells compared to CD20^−^ NK cells (Figure 3, F to H). There was no significant difference in the ratio of granulysin expressing cells between the two groups (Figure 3I). For GrK, the ratio was higher among CD56^+^CD20^+^ cells (Figure 3J). These results are in line with our data showing that CD20^+^ NK cells are relatively more common among CD56^bright^ NK cells. Of note, further analysis revealed that the GrK^+^ cell ratio was similar and close to 100% in both CD56^bright^CD20^+^ and CD56^bright^CD20^−^ cells (Figure S3, A and C), whereas a higher ratio CD56^dim^CD20^+^ cells expressed GrK compared to CD56^dim^CD20^−^ cells (Figure S3, B and D).

We also found that a greater proportion of the CD56^+^CD20^+^ cells express death inducing ligands FasL, and TRAIL on the cell surface (Figure 3K).

In line with the above findings, CD56^+^CD20^+^ cells secreted higher amounts of GrA, GrB, perforin, soluble FasL and soluble Fas into the cell culture medium after cytokine stimulation (Figure 3L).

### CD20^+^ NK cells have a higher capacity of degranulation and are more effective at killing K562 leukemia cells

To investigate the functional consequences of these findings, we first performed a CD107a degranulation assay. We discovered that the percentage of degranulating cells was two times higher among CD20^+^ NK cells compared to CD20^−^ NK cells after incubation with K562 cells (median, IQR: 37.1, 30.1 – 41.2 vs 16.6, 13 – 19) (Figure 4A). Furthermore, the degranulation capacity of individual CD20^+^ NK cells was found to be significantly higher as revealed by MFI of the cell surface CD107a molecules measured after stimulation (median, IQR: 25636, 23568 – 29864 vs 19655, 17674 – 21876) (Figure 4B).

**Fig. 4.**
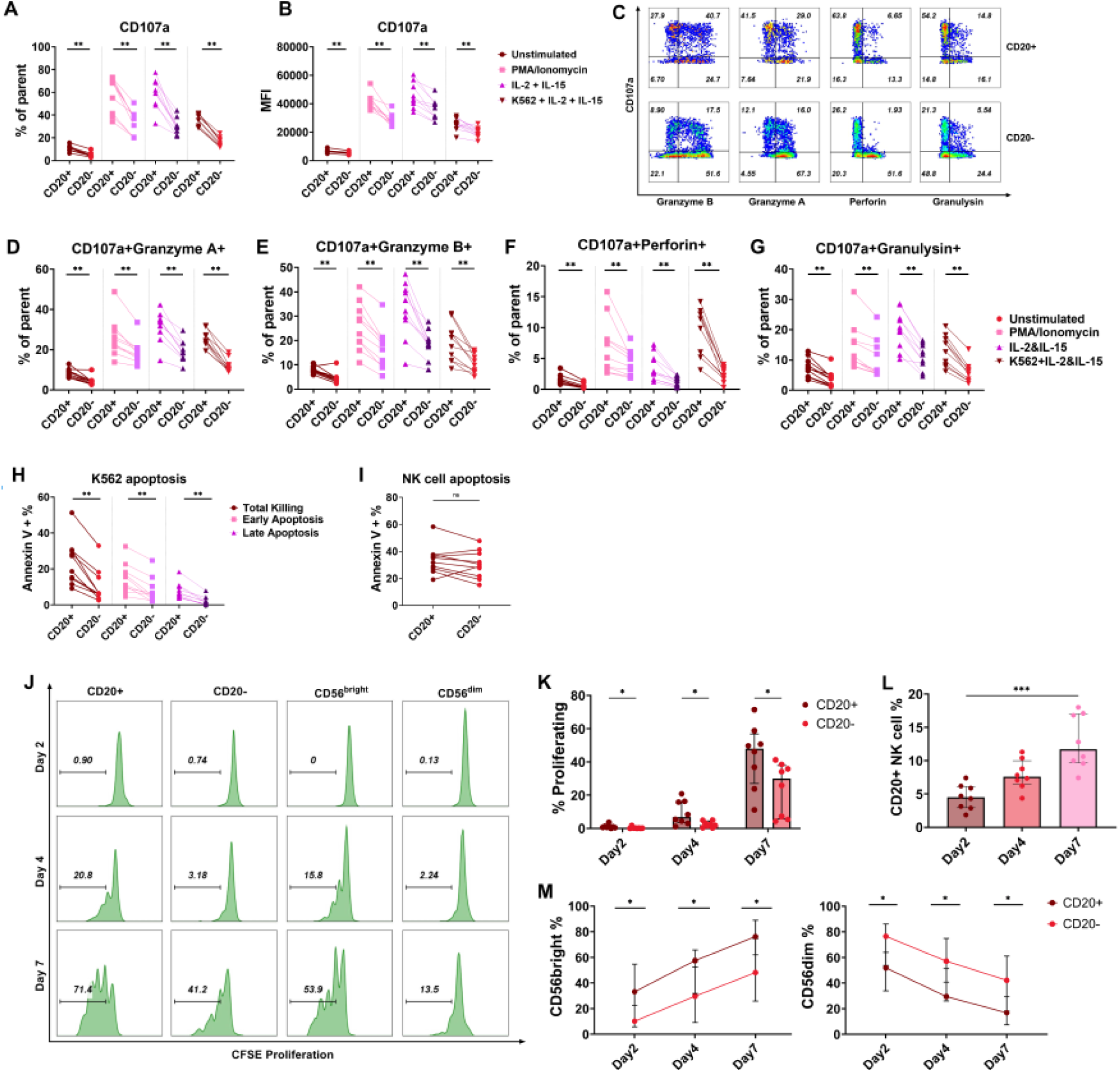
CD20^+^ NK cells are more efficient killers and have a higher capacity for proliferation after stimulation. **(A)** A higher percentage of CD20^+^ NK cells degranulate after stimulation and **(B)** degranulation capacity of individual CD20^+^ NK cells are higher. **(C)** Representative intracellular staining of degranulating CD20^+^ and CD20^−^ NK cells. **(D to G)** Degranulating CD20^+^ NK cells have a higher content of GrA (C), GrB (D), perforin (E) and granulysin (F) compared to CD20^−^ NK cells. **(H)** CD20^+^ NK cells are more efficient killers compared to CD20-NK cells. P values were calculated using Wilcoxon matched-pairs signed rank test. **(I)** The rate of apoptosis is similar between CD20^+^ and CD20^−^ NK cells after killing. **(J)** Representative flow cytometry analysis of CFSE stained NK cell subgroups. **(K)** CD20^+^ NK cells have a higher proliferative capacity under IL-2 and IL-15 stimulation. P values were calculated using Wilcoxon matched-pairs signed rank test with Holm-Sidak’s post hoc analysis. **(L)** Accordingly, CD20^+^ NK cell ratio increased gradually after stimulation. P values were calculated using Friedman test with Dunn’s multiple comparisons post hoc correction. **(M)** CD56^bright^ cells can proliferate more compared to CD56^dim^ cells in both CD20^+^ and CD20^−^ populations. *P < 0.05, **P < 0.01, and ***P < 0.001.

We also analyzed the frequency of GrA, GrB, perforin and granulysin expressing cells among degranulated cells. We found that degranulating CD20^+^ NK cells contain a higher ratio of all four cytotoxic molecules compared to CD20^−^ NK cells (Figure 4, C to G). These findings show that, CD20 expression is associated with superior degranulation capacity for cytotoxic molecules in NK cells.

Next, we performed a killing assay with K562 cells co-cultured with either sorted CD20^+^ or CD20^−^ NK cells to directly compare their cytotoxic efficacy. Indeed, CD20^+^ NK cells were found to be more efficient killers compared to CD20^−^ NK cells (p<0.01) (Figure 4H). We also measured the ratio of NK cells that undergo apoptosis to see if CD20^+^ NK cells are more prone to cell death after activation. We did not detect any difference between CD20^+^ and CD20^−^ NK cells (Figure 4I).

### CD20^+^ NK cells have a higher proliferative capacity under IL-2 and IL-15 stimulation

Finally, we checked whether CD20^+^ NK cells can be expanded with long term cytokine stimulation by incubating carboxyfluorescein succinimidyl ester (CFSE) stained PBMCs from healthy volunteers with IL-2 and IL-15 for seven days. We found that CD20^+^ NK cells have a higher proliferative capacity in response to cytokine stimulation (p<0.05 at day 2, 4 and 7) (Figure 4J and K). Moreover, the CD20^+^ NK cell ratio increased gradually (p<0.001, Figure 4L). As expected, CD56^bright^ cells proliferated more in both CD20^+^ and CD20^−^ cell groups (Figure 4M).

### Molecular features of CD56^+^CD20^+^ cells

So far, we have identified that CD56^+^CD20^+^ cells are highly functional cells with increased proliferative capacity and are expanded in neurological disorders, especially MS. Increased functionality and proliferative capacity are characteristic features of memory-like NK cells. To gain more insight into the molecular underpinnings of these characteristics, we analyzed selected publicly available scRNAseq datasets generated from patients with inflammatory conditions.

First, we analyzed data from Twin-MS study, in which frozen PBMC obtained from eight MS and non-MS monozygotic twin pairs and two healthy samples were used (*11*). This dataset was created by single-cell RNA sequencing of sorted CD3^+^CD4^+^ and CD3^−^CD11c^+^ cells. The dataset contains 56801 NK cells and 1614 (2.84%) of these cells had *MS4A1* expression, the gene coding for CD20. The percentage of *MS4A1* expressing NK cells was 3.2% in twins with MS and 2.6% in twins without MS. The percentage of *MS4A1* expressing cells was highest in the CD56^bright^-like cluster that is characterized by high *GZMK* expression (7.44%), followed by proliferating NK cell (2.96%) and CD56^dim^-like (1.39%) clusters (Figure 5, A to E). The analysis of differentially expressed genes (DEGs) between NK cell subclusters verified the presence of *MS4A1* among the top upregulated genes in the CD56^bright^-like cluster (table S2).

**Figure 5.**
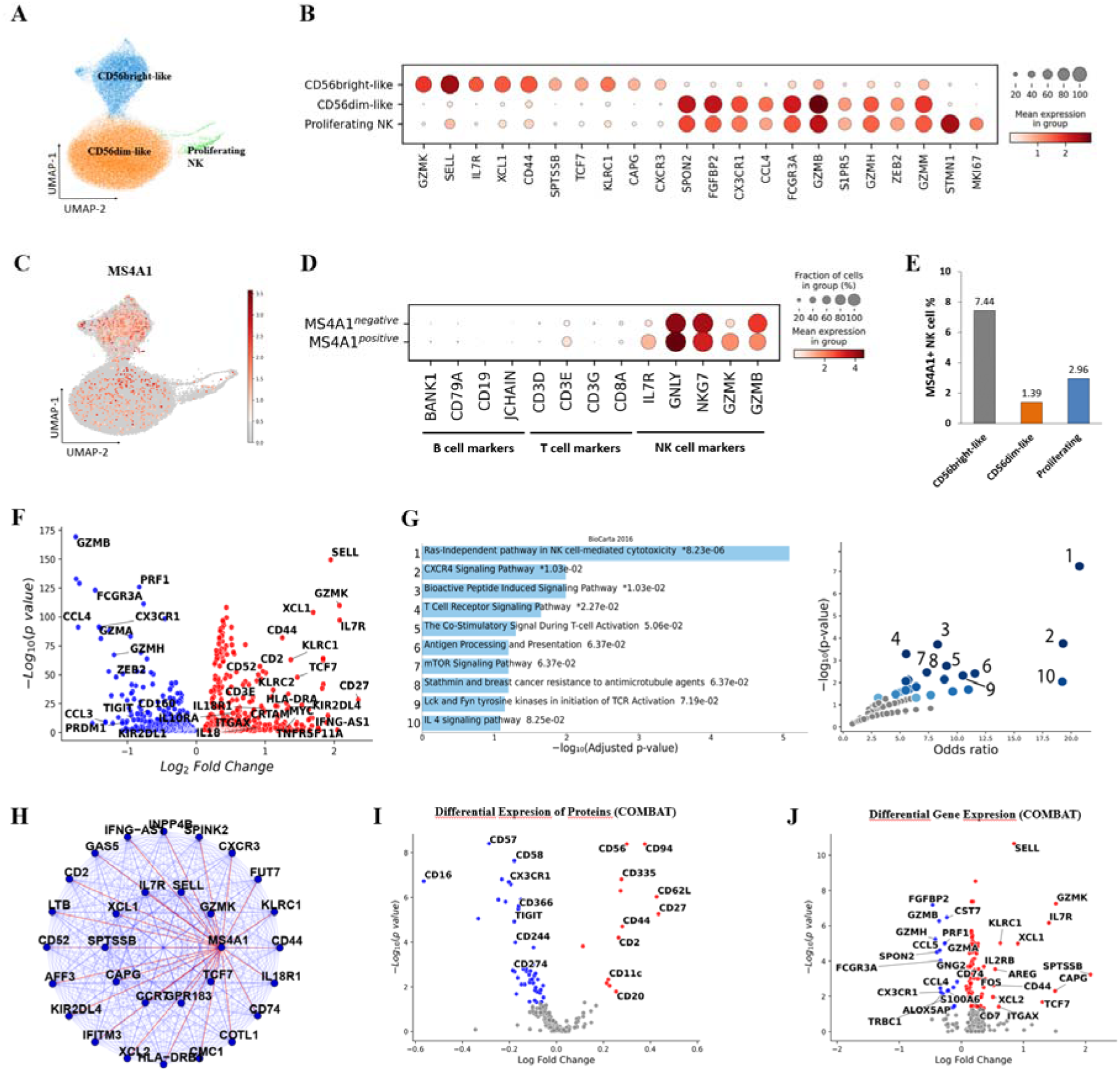
Molecular features of *MS4A1* expressing NK cells in the Twin-MS dataset. **(A)** UMAP graph showing the identified NK cell subclusters. **(B)** The dot plot of the top differentially expressed genes in each subcluster. **(C)** UMAP plot demonstrating the distribution of *MS4A1*-expressing NK cells (red dots, enlarged for clarity). **(D)** Dot plot showing expression of marker genes for B cells, T cells and NK cells demonstrates that there is no B or T cell contamination in the NK cell cluster. **(E)** *MS4A1*-expressing NK cells are enriched in the CD56^bright-like^ subset. **(F)** Volcano plot demonstrating the differentially expressed genes in *MS4A1*-expressing NK cells versus others. **(G)** The pathways that are active in *MS4A1* expressing NK cells revealed by the geneset enrichment analysis of the upregulated genes by using the BioCarta 2016 database (left). The graph showing the p-value and odds ratio of each pathway (right). **(H)** The network of genes co-expressed with *MS4A1* in the CD56^bright^-like module, as revealed by WGCNA. Lines connecting the dots represent connectivity between the genes.

Next, we performed differential gene expression (DGE) analysis of *MS4A1* expressing NK cells compared to other NK cells. We found that expression of *GZMK*, in addition to the genes encoding for cytokine receptors (*IL7R, IL18R1, IL18RAP, IL4R, IL10R, IL12RB2, IL2RB, IL2RG*), NK cell receptors (*KLRC1, KLRC2, KLRC3, KLRC4, KLRD1, KLRK1, KIR2DL4*), TNF receptors (*CD27, TNFRSF11A, TNFRSF18*), and NK cell memory related surface markers (*SELL, CD44, CD2, CD52, CD3E, HLA-DR*) are increased in *MS4A1* expressing NK cells (Figure 5F, table S3). On the other hand, the genes belonging to mature NK cells such as *TBX21, PRDM1, GZMA, GZMB, PRF1, FCGR3A, NKG7, KLRG1, CX3CR1, S1PR5, CCL3, CCL4, CCL5,* and the genes coding for inhibitory receptors such as *TIGIT, HAVCR3, CD160 and inhibitory KIR receptors* were downregulated (Figure 5F, table S3). An analysis of gene ontology for the upregulated genes in MS4A1-expressing NK cells unveiled several significant biological pathways. These pathways encompassed cytoplasmic translation machinery, leukocyte cytotoxicity, assembly of MHC class II protein complexes, cell adhesion, and intracellular signal transduction molecules (Table S4). Gene set enrichment analysis of the upregulated genes in MS4A1-expressing NK cells indicated the activation of pathways associated with RAS-independent NK cell-mediated cytotoxicity and signaling induced by bioactive peptides (Figure 5G, table S5).

Finally, we conducted weighted correlation network analysis (WGCNA) on NK cells within the Twin-MS dataset to test whether *MS4A1* appears in any transcriptional module. We found that *MS4A1* is transcribed within the module distinguished by high GZMK expression, confirming higher expression of *MS4A1* in the CD56^bright^-like cluster. *MS4A1* expression correlated positively with genes including *SELL, IL7R, GZMK, XCL1 and TCF7* (Figure 5H, table S6).

Next, we analyzed the COMBAT study dataset, a CITE-seq multi-omics dataset derived from the PBMC samples of COVID-19 patients (*12*). This dataset contained 69927 NK cells, 884 (1.26%) of which expressed *MS4A1*. Importantly, DGE analysis of cell surface proteins validated that CD20 protein expression is higher on *MS4A1*-expressing NK cells compared to other NK cells (Figure 5I, table S7). Moreover, CD56, CD27, CD62L, CD44, NKp46 (CD335), CD2, and CD11c were more abundant, while CD16, KLRG1, CD57, and CX3CR1 were less abundant on these cells, confirming our results obtained by flow cytometry analysis (Figure 5I, table S8). Additionally, *MS4A1* expressing cells were again more frequent in the CD56^bright^ subcluster (Figure 5J), and the DGE analysis gave a list of upregulated genes that highly overlap with the results of the Twin-MS dataset (Figure 5J, table S8).

The results so far show that CD20^+^ NK cells are enriched in the CD56^bright^ cell population, which is predominantly present in the tissues, especially secondary lymphoid organs, and underrepresented in blood (*13*). Therefore, to get a more detailed picture of the *MS4A1* expressing NK cells, we analyzed another scRNAseq dataset that combines data from FACS-sorted CD45^+^CD56^+^CD3^-^CD14^-^CD15^-^CD163^-^ NK cells of paired blood and resected metastatic melanoma tissues obtained from five patients (*14*). *MS4A1* expressing NK cell ratio was 0.35% (69/19859) in blood, whereas it was much higher (4.21%, 545/12936) in tumor tissue (Figure 6, A to C). The percentage of *MS4A1* expressing NK cells was again highest among the *IL7R^+^GZMK^+^GZMB*^low^ CD56^bright-like^ cells (cluster 1, 6%), followed by *IL7R^+^GZMK^−^GZMB*^−^ NK precursor (NKP) cells (cluster 3, 3.4%), *STMN1^+^* proliferating NK cells (cluster 4, 1.8%), *IL7R^−^ GZMK^+^GZMB*^low^ CD56^int^ cells (cluster 2, 1.15%) and *IL7R^−^GZMK^−^GZMB*^high^ CD56^dim-like^ cells (cluster 0 and 5; 0.06% and 0%, respectively) (Figure 6, D to H). The DGE analysis of *MS4A1* expressing NK cells yielded highly overlapping results with the Twin-MS and COVID datasets in terms of upregulation of the CD56^bright^ related genes and downregulation of the CD56^dim^ related genes in both blood and tumor (Figure S4, A and B; tables S9 and S10, respectively).

**Figure 6.**
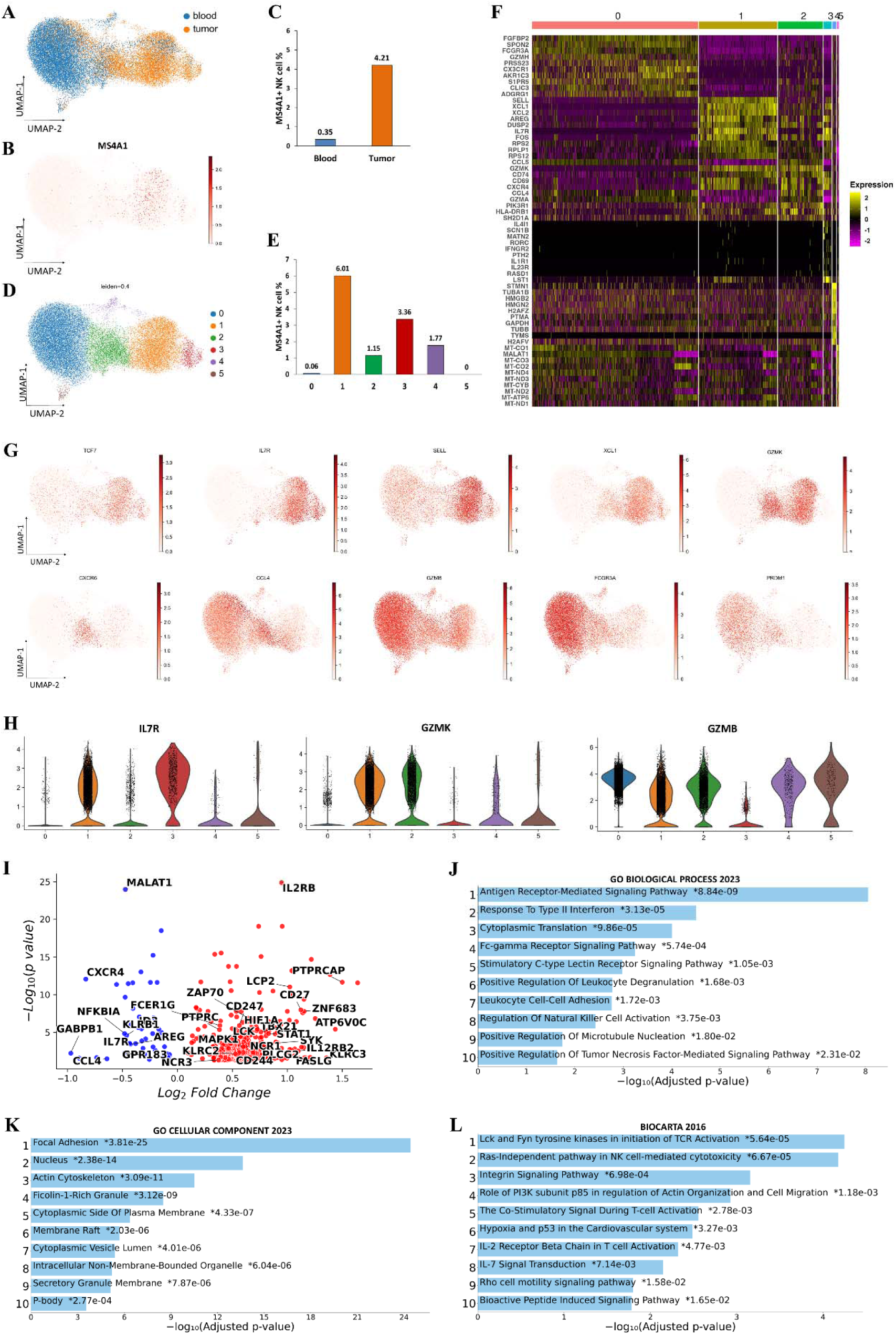
Distribution and molecular features of *MS4A1* expressing NK cells in the paired blood and tumor samples from patients with melanoma. **(A)** UMAP plot shows that NK cells obtained from the blood and tumor tissues cluster in opposite sides and have little overlap. **(B and C)** *MS4A1* expression is scarce in blood NK cells whereas it is dramatically enriched in tumor tissues. **(D)** UMAP graph showing the major subclusters of NK cells (Leiden 0.4). (E) *MS4A1* expression is highest in cluster 1, followed by clusters 3 and 4; whereas it is very low in clusters 0 and 5. **(F)** Heatmap demonstrating the highly expressed genes in each cluster. **(G)** Selected genes that are enriched in various clusters are highlighted in the UMAP plot. *TCF7, IL7R, SELL* and *XCL1* are highly expressed in both cluster 1 (CD56^bright-like^) and 3 (NKP cluster), whereas *GZMK* is expressed by the clusters 1 and 2 (CD56^int^), but not 3. *CXCR6* and highly *CCL4* are enriched in cluster 2. *GZMB, FCGR3A* and *PRDM1* are highly expressed by the CD56^dim-like^ clusters 0 and 5. **(H)** Bar plots showing the gene expression level of marker genes *IL7R, GZMK* and *GZMB* in different NK subclusters. **(I)** Volcano plot showing the DGEs in *MS4A1* expressing NK cells of cluster 1 (CD56^bright-like^). **(J and K)** GO annotation analysis of the upregulated genes in *MS4A1* expressing NK cells of cluster 1. **(L)** GSEA analysis of *MS4A1* expressing NK cells of cluster 1 according to the BioCarta 2016 database.

Next, to gain more insight into the molecular mechanisms behind the activated state of *MS4A1*^+^ NK cells, we conducted a DGE analysis restricted to the cells in cluster 1, that includes 83% (511/614) of all *MS4A1* expressing NK cells. This analysis yielded 561 upregulated and 51 downregulated genes in *MS4A1* expressing *IL7R^+^GZMK^+^GZMB*^low^ CD56^bright-like^ cells (Figure 6I, table S11). Gene ontology analysis of the upregulated genes, based on cellular components, indicated associations with focal adhesions, the nucleus, the actin cytoskeleton, and lipid rafts. Analysis of biological processes revealed involvement in receptor-mediated signaling (C-type lectin receptor and Fc-γ receptor pathways), responses to type II IFN and TNF, leukocyte degranulation, NK cell activation, cell adhesion, protein translation, and microtubule nucleation (Figure 6, J and K; Figure S4, C and D; tables S12 and S13). Gene set enrichment analysis showed activation of the pathways related to receptor signaling through Lck and Fyn tyrosine kinases, Ras-independent NK cell-mediated toxicity (through NKG2C and NKG2E receptors), actin organization, cell migration, hypoxia response, and STAT1 signaling. (Figure 6L, Figure S4E, table S14). Additionally, CD27, IL12RB2, IL2RB, TBX21, CD244, NCR1, NCR3, CD247 and FCER1G were also upregulated highlighting the highly responsive status of CD20^+^ NK cells to IL12, IL15 and activating NK cell receptors, and tendency of these cells to differentiate into a mature type 1 immune cell phenotype upon stimulation.

In summary, these results provide a comprehensive explanation for the molecular basis of the remarkably active cell state observed in CD20^+^ NK cells, as revealed by the flow cytometry experiments, despite their seemingly less mature phenotype.

### CD56^+^CD20^+^ cells are expanded in inflammatory disorders, including chronic HBV infection, hepatocellular carcinoma, autoimmune hepatitis, and lung cancer

Our results so far indicate that CD56^+^CD20^+^ cells display features of memory-like NK cells. Therefore, we wondered whether expansion of CD56^+^CD20^+^ cells is seen in other inflammatory disorders. First, we analyzed paired peripheral blood and hepatic tissue from patients with alcohol related liver disease and chronic hepatitis B virus (HBV) infection with or without hepatocellular carcinoma (HCC) (table S15). CD56^+^CD20^+^ cells were expanded in the blood and liver tissues obtained from patients with HBV^+^ HCC patients compared to the patients only with chronic HBV infection (Figure 7, A and B). Moreover, there was a tendency for CD56^+^CD20^+^ cell expansion in persistent HBV infection -related chronic liver disease compared to alcohol-related liver disease, again in both tissue types tested (Figure 7, A and B). In addition, CD56^+^CD20^+^ cells were enriched in the liver compared to the paired blood samples (Figure 7C).

**Figure 7.**
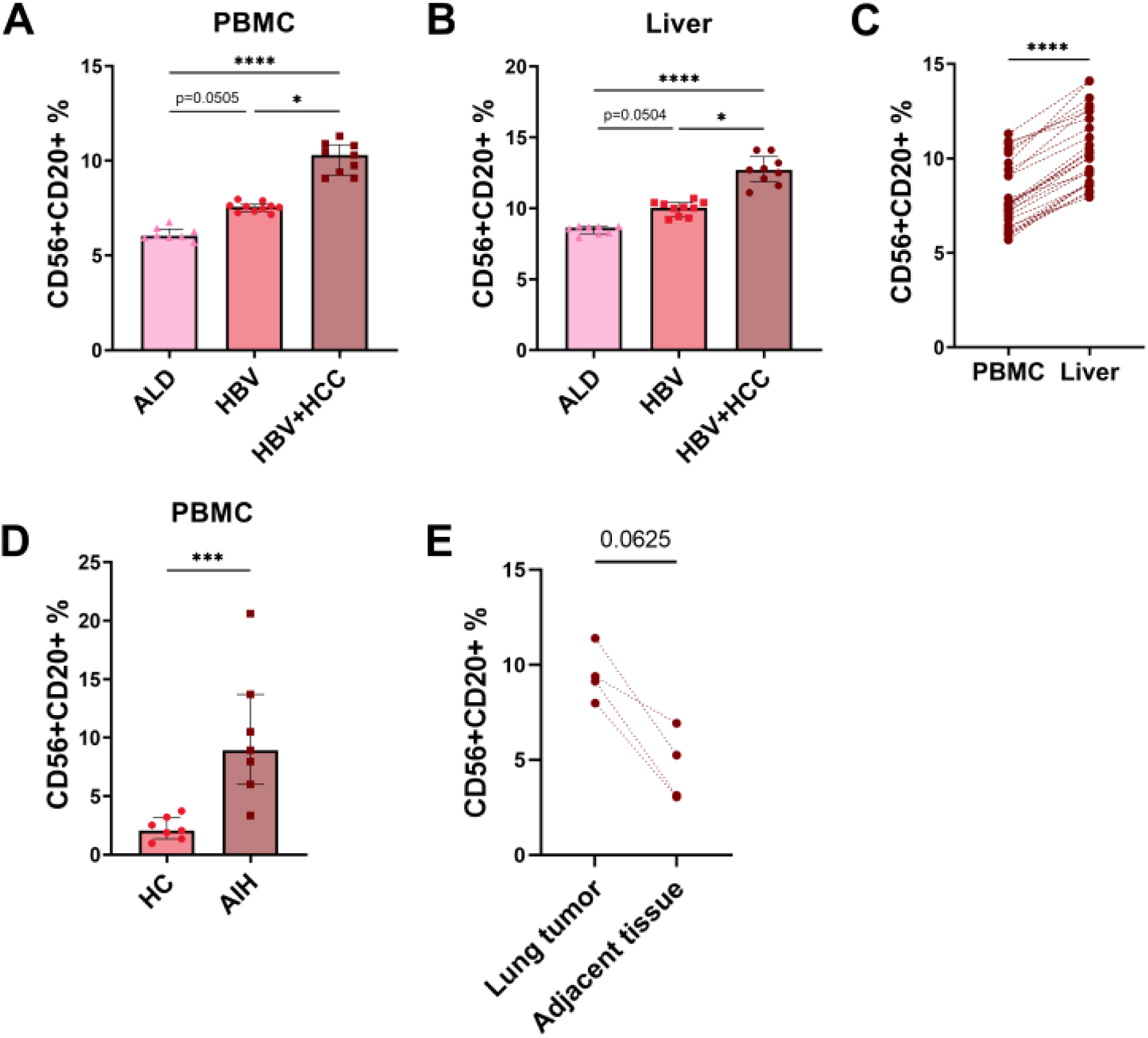
CD20^+^ NK cells are increased in the tissues of patients with inflammatory disorders. **(A and B)** CD56^+^CD20^+^ cells were expanded in the blood (A) and liver (B) tissues obtained from patients with HBV-related chronic liver disease and hepatocellular carcinoma (HBV^+^ HCC) (n=9) compared to the patients only with chronic HBV (n=10) infection. In addition, there was a strong tendency for CD56^+^CD20^+^ cell expansion in HBV-related chronic liver disease compared to alcoholic liver disease (ALD) (n=8), again in both tissue types (A and B). P values were calculated using Kruskal-Wallis test with Dunn’s multiple comparisons post hoc correction. **(C)** CD56^+^CD20^+^ cells are enriched in the liver of patients with chronic liver diseases (ALD, HBV and HBV^+^ HCC) compared to the paired peripheral blood (n=27). **(D)** The ratio of CD56^+^CD20^+^ cells are increased in the blood of children with autoimmune hepatitis (AIH) (n=7) compared to healthy controls (HC) (n=7). **(E)** There was a tendency for the enrichment of CD56^+^CD20^+^ cells inside the lung tumor compared to the paired surrounding tissue (n=4). P values were calculated using Wilcoxon matched pairs signed rank test.

Next, we investigated children with autoimmune hepatitis and found that CD56^+^CD20^+^ cells are more frequent in the peripheral blood of patients compared to HCs (Figure 7D, table S15).

Lastly, we examined samples of surgically removed lung cancer tissue and neighboring healthy tissue. Again, there was a tendency for the enrichment of CD56^+^CD20^+^ cells in the cancer tissue compared to the neighboring tissue (Figure 7E).

### *MS4A1* expressing NK cells are enriched in secondary lymphoid organs

Our results show that CD56^+^CD20^+^ cells carry a lymph node homing capacity, characterized by high expression of *SELL*. Therefore, we investigated the distribution of CD56^+^CD20^+^ cells throughout the body in the publicly available human cross-tissue immune cell analysis dataset by Dominguez Conde et al. (*15*).

This is an adult tissue dataset and combines independent data from two sources: curated information from published studies and original data generated by the study team. First, we observed that *MS4A1* is expressed in a small fraction of NK cells in the bone marrow in both resources. Second, we observed that the ratio of *MS4A1* expressing NK cells varied among tissues. The percentage was relatively lower in the peripheral blood, bone marrow and most solid tissues, whereas it was higher in the secondary lymphoid organs, especially in the spleen. The rate of CD20 expression was highest among the CD56^bright^-like NK cluster characterized by high *GZMK* and *SELL* expression, in line with our previous findings (Figure 8A, table S16). By using flow cytometry, we verified the presence of CD20^+^ NK cells in the recently transplanted bone marrow obtained from patients with leukemia who were treated with immune suppression followed by bone marrow transplantation (Figure 8B).

**Figure 8.**
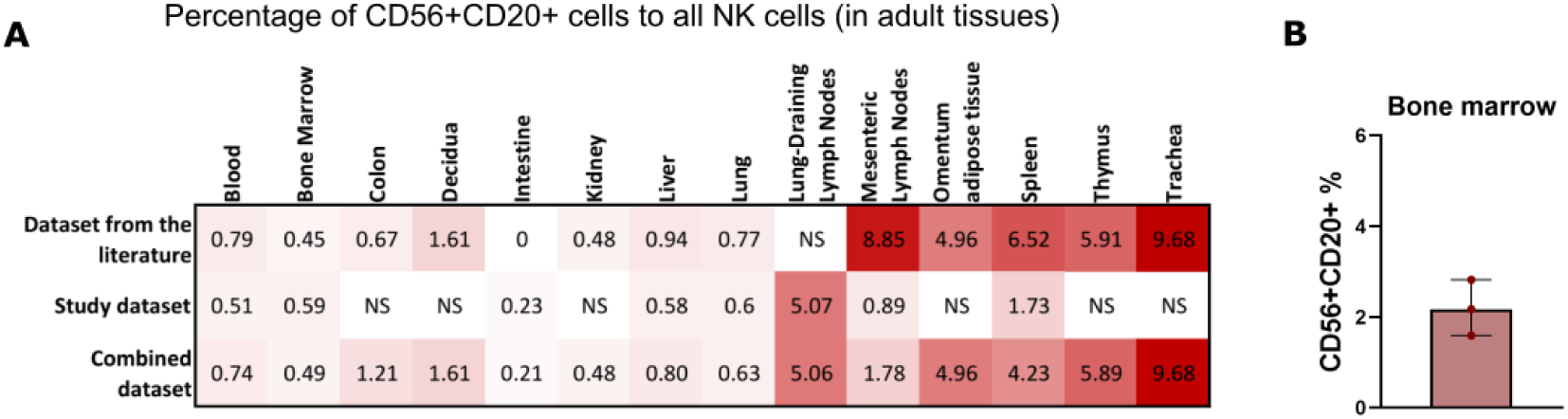
CD20^+^ NK cells are abundant in secondary lymphoid organs, and present during development. **(A)** Heatmap showing the distribution of *MS4A1* expressing NK cells in different organs in the cross-tissue immune cell analysis study which was performed by the analysis of adult human tissues. Enrichment of these cells in the secondary lymphoid organs is evident. Numbers show the percentage of *MS4A1* expressing NK cells in each organ. Data from tissues with less than 100 available cells were not shown (NS). **(B)** The presence of CD20^+^ NK cells in the bone marrow was verified by flow cytometry in patients with leukemia and recent bone marrow transplantation.

## DISCUSSION

Herein, we identified that a fraction of NK cells expresses CD20. These CD56^+^CD20^+^ cells have an active phenotype and superior functional capacity, in terms of cytotoxicity and cytokine secretion. Despite increased functionality, these cells are not terminally differentiated and carry homeostatic and memory-like features, evidenced by high *TCF7*, *IL7R, CD27* expression and higher proliferative capacity. CD56^+^CD20^+^ cells are preferentially located in secondary lymphoid organs and are expanded in the blood and disease-relevant tissues in inflammatory disorders.

The features of CD20^+^ NK cells identified in this study exhibit similarities to CD20^+^ T cells. CD3^+^CD20^+^ cells are recognized as proinflammatory cells, characterized by an activated phenotype and increased cytokine secretion upon stimulation (*4, 5, 8*). Likewise, our findings indicate an elevated proportion of CD20^+^ NK cells expressing the activation markers CD69 and CD137, along with the activating receptor NKp46. Additionally, we observed heightened production and secretion of TNF-α, IFN-γ, GM-CSF, and IL-10 after stimulation in CD56^+^CD20^+^ cells. Phenotypically, CD3^+^CD20^+^ cells are predominantly enriched in Th1 and Tc1 memory subsets, exhibiting lower abundance among naïve and terminal effector subsets (*5*). Similarly, we observed a diminished proportion of CD56^+^CD20^+^ cells expressing markers associated with mature and terminally differentiated NK cells, such as CD57, CD16, GzB, and perforin. A striking feature of CD20^+^ NK cells is their heightened cytotoxic activity. While previous research has shown that CD3^+^CD20^+^ cells are enriched for CD8^+^ cytotoxic T cells, it remains uncertain whether these cells also exhibit elevated cytotoxicity, necessitating further investigation in future studies.

Our investigation revealed a higher percentage of CD20 expression among CD56^bright^ cells as compared to CD56dim NK cells. CD56^bright^ NK cells are known for their potent cytokine-producing abilities with restricted cytotoxicity. However, we found that CD20^+^ NK cells exhibit greater cytotoxicity and efficiency in killing K562 cells. Several findings reveal the cellular mechanisms behind the superior cytolytic potential of CD56^+^CD20^+^ cells. First, CD56^+^CD20^+^ cells exhibit enhanced degranulation, with higher granule secretion per cell and increased expression of cytotoxic molecules compared to CD20^−^ cells. Second, CD56^+^CD20^+^ cells display elevated levels of apoptosis-inducing receptors FASL and TRAIL on their surface. Collectively, these features underscore the enriched cytotoxic arsenal of CD20^+^ NK cells, coupled with an efficient molecular machinery facilitating effective lytic granule release. These observations strongly indicate a primed state in these cells, poised for robust type 1 immune responses.

Analysis of publicly available scRNAseq data unveiled the molecular basis underlying this heightened polyfunctionality. We observed upregulation of ITAM-bearing NK receptor complexes *CD247* (CD3ζ) and *FCER1G* (Fc RI-γ), along with members of Src family kinases (*LCK, LYN, FYN, FGR*), tyrosine kinases (*SYK* and *ZAP70*), *PLCG2, LCP2,* and several actin cytoskeleton and membrane raft-related molecules like *ZYX, LCP1, ACTB RFTN1,* among others, in *MS4A1* expressing NK cells. Additionally, genes associated with cytolytic granule formation and microtubule organization were also found to be upregulated in *MS4A1* expressing NK cells. These findings suggest that CD20 may interact with natural killer cell activating receptors, leading to an amplification of stimulatory signals upon stimulation. This, in turn, may trigger protein synthesis, cytoskeleton reorganization, ultimately culminating in increased cytokine and cytotoxic granule release. Identification of the specific protein-protein interactions will require further studies.

We found that expression of *MS4A1* correlates with several other genes including *TCF7, GZMK, SELL, IL7R and XCL1*. *TCF7* codes for the transcription factor Tcf1 that plays a critical role in NK cell survival during maturation and upon activation by limiting excessive granzyme B expression (*16*). GzK is known for its prominent pro-inflammatory function and its role in endothelial activation and breaking of the blood-brain barrier (*17*). Recently, GrK expressing CD8^+^ T cells have been described as tissue-enriched cells and reported to constitute the core inflammatory population in tissues undergoing inflammation (*18*). We observed a dramatic upregulation of GrK in CD20^+^ NK cells after incubation with IL2, IL15 and K562 cells. Thus, GrK expression may be a significant contributor of the tissue-enriched and pro-inflammatory nature of CD56^+^CD20^+^ cells.

Molecules previously linked to enhanced functionality and memory-like features in NK cells are similarly upregulated in CD20+ NK cells. Among these, *SELL* encodes for CD62L. CD62L^+^ CD56^dim^ NK cells were previously reported to be polyfunctional cells with a higher killing and cytokine production capacity and preserve their ability to proliferate during viral infection (*19*). *CD44* has a role in cytoskeleton rearrangement and raft reorganization and signaling through CD44 has been shown to enhance NK cell cytotoxic function. CD2 is important for NK cell immune synapse formation and recruits CD16 to the immune synapse, lowering the threshold for activation. It is upregulated in adaptive NK cells even in the absence of NKG2C (*20*). Finally, *CD52* was reported among the top upregulated genes together with *KLRC2*, *CD2* and *CD3E* in the adaptive NK cell population in CMV seropositive patients in two recent studies (*21, 22*). Another memory-like feature of CD56^+^CD20^+^ cells is their higher proliferative capacity in response to IL-2 and IL-15 stimulation.

Our investigation encompassing two distinct cohorts of individuals affected by neurological and hepatic disorders revealed an expansion of CD56^+^CD20^+^ cells in both the blood and disease-relevant tissues. Remarkably, the degree of expansion was more pronounced in MS compared to other neurological disorders, and it was also higher in individuals with HBV^+^ HCC compared to those with chronic HBV-related chronic liver disease. Additionally, we observed a strong correlation between CD20^+^ NK cells and CD20^+^ T cells and B cells in individuals with MS, highlighting the relevance to these cells in adaptive immune response. Taken together, these clinical findings, along with the molecular insights provided above, suggest that elevated CD20 expression may be a common mechanism in various lymphocyte subtypes during chronic inflammatory conditions and is associated with enhanced functionality and memory features.

There are some limitations of our study. First, while our flow cytometry and functional assays identified distinctive features of CD20+ NK cells, these findings were primarily based on in vitro assays, which may not fully capture the in vivo behavior of these cells in complex tissue environments. Second, although we analyzed publicly available single-cell RNA sequencing datasets to validate our findings, the limited availability of high-resolution datasets from diverse patient populations may restrict the generalizability of our results across different inflammatory and oncologic conditions. Third, while we observed an enrichment of CD20+ NK cells in inflamed tissues and peripheral blood of patients with various inflammatory diseases, the underlying mechanisms driving their expansion and precise role in disease progression remain to be fully elucidated. Finally, our study did not address the long-term stability and functional plasticity of CD20+ NK cells, which may be relevant for understanding their potential as therapeutic targets. Future studies should aim to address these limitations by examining CD20+ NK cells in more diverse patient cohorts and exploring their *in vivo* functions in disease models. In conclusion, we have identified that CD20 expression is not restricted to B cells and T cells and NK cells can also express CD20. These CD56^+^CD20^+^ NK cells exhibit enhanced cytokine production, cytotoxicity, and proliferative capacity, and are expanded in various inflammatory disorders. The unique features of CD56^+^CD20^+^ cells render them both as possible treatment targets and as therapeutic tools. Further investigations are warranted to investigate the role of CD56^+^CD20^+^ cells in various disorders.

## MATERIALS AND METHODS

### Study Population

Neurological cohort of the study was approved by the Koç University Clinical Research Ethics Committee under protocol number 2020.456.IRB1.169. A voluntary informed consent form was signed by all patients and healthy controls. 31 MS, 8 IDD, 12 OIND and 15 NINS patients and 14 healthy volunteers were included in the study. All patients received their diagnosis from Bakırköy Mazhar Osman Psychiatric Hospital Neurology Outpatient Clinics depending on EDSS scores, Oligoclonal Band Electrophoresis results and FMRI scans. 10ml of Peripheral blood were drawn into vacutainer tubes containing K_3_EDTA and CSF were drawn into non-coagulated tubes by lumbar puncture. Samples were transported to Koç University Hospital Research Center for Translational Medicine. Blood and CSF samples of MS and IDD patients were analyzed as patient population whereas OIND and NINS patients’ blood samples were analyzed as disease controls. EDSS scores of MS patients were also collected to be compared with CD20^+^ NK cell frequencies in both blood and CSF samples.

To determine if CD56^+^CD20^+^ cells are really a subtype of NK cells, we analyzed PBMCs of 12 MS patients before and after 6 months of Rituximab treatment. Samples were obtained from Ludwig Maximillan University Institute of Clinical Neuroimmunology and analyzed at LMU Flow Cytometry core facility using LSR Fortessa (BD Biosciences, USA)

For the chronic liver disease (CLD) cohort, tissue collection and liver mononuclear cell isolation methods adhered to ethical guidelines. 27 CLD patients were enrolled in this study, with approval from the Koç University Ethics Committee under approval numbers 2016.024.IRB2.005 and 2017.139.IRB2.048. Liver specimens were collected from explant livers, and histopathological scoring followed the Ishak pathological staging system. The study population included patients with hepatocellular carcinoma alongside viral hepatitis infection, alcohol-related liver disease, or viral hepatitis. Male and female participants with alcohol consumption exceeding 60 g and 40 g per day, respectively, were included in the alcohol-related liver disease cohort. BCLC stage for HCC and FIB-4 scores for all CLD were assessed as part of the analysis.

To assess the presence of CD20 expressing NK cells in autoimmune disorders, 7 pediatric patients with autoimmune hepatitis and 7 age and sex matched controls were also included into the study with approval from the Koç University Ethics Committee under approval number 2019.255.IRB2.077. Liver biopsies of healthy controls were also analyzed to determine CD20^+^ NK cells in liver. Frozen PBMC samples were acquired from Koç University Hospital Department of Pediatric Hepatology.

Tumor and adjacent healthy tissues from 4 lung cancer patients at Koç University Hospital were analyzed also to observe if CD20^+^ NK cells were abundant in cancer tissues. For this purpose, tissue samples were homogenized into single cell suspensions by GentleMACS Octo Dissociator using Human Tumor Dissociation Kit (Milteny Biotech, Germany). Patients were enrolled in the study with approval from the Koç University Ethics Committee under approval number 2020.001.IRB2.001.

Apart from the Rituximab treated patient samples, every other sample were analyzed at Koç University Hospital Research Center for Translational Medicine (KUTTAM) Flow Cytometry core facility using Attune NxT Focusing Cytometer (Thermo Scientific, USA). Flow Cytometry data were stored as FCS files and analyzed by FlowJo v.10.10.0 (BD Biosciences, USA)

### PBMC Isolation

PBMCs from peripheral blood were isolated using density gradient centrifugation method. Blood samples were diluted 1:1 with staining buffer (PBS containing 1% BSA), carefully layered on equal volume of Lymphoprep d=1.077g/ml (Axis-Shield, Norway) and centrifuged at 500g for 30 minutes at room temperature without brakes. Buffy coat containing PBMCs were collected into a sterile 50ml falcon tube and washed with equal volume of staining buffer at 500g for 5 minutes. Supernatant was discarded and if red blood cells were observed, cell pellet was resuspended with 2 ml of red blood cell lysis buffer (NH_4_Cl, Na_2_HCO_3_, EDTA, pH = 7.3), incubated for 10 minutes at room temperature and centrifuged at 500g for 5 minutes. After centrifugation, cell pellet was resuspended with 1 ml of staining buffer and counted on Sysmex XE-2100 Automated Cell Counter (Roche, Germany). 1×10^7^ cells were cryopreserved in liquid nitrogen whereas 6×10^6^ cells were used in immunophenotyping experiments.

### Liver Mononuclear Cell Isolation

Mononuclear cells from explanted livers were isolated using a non-enzymatic method paired with Ficoll gradient separation. The liver tissue was finely minced into smaller fragments. Serum-free RPMI-1640 medium was added to a stomacher bag containing the minced tissue, which was then incubated in a Stomacher machine (Seward Laboratory Systems, UK) to facilitate cell release. After incubation, the cell mixture was extracted from the stomacher bag and filtered through a 125 μm nylon mesh. The filtrate was subjected to washing steps, and the resulting pellet was resuspended in 1X DPBS (Thermo Scientific, USA). The cell suspension was gently layered over Lymphoprep (density: 1.077 g/mL, Axis-Shield, Norway) to create a distinct separation layer followed by centrifugation. An additional washing step was performed to remove residual Lymphoprep. Finally, the pellet was resuspended in a freezing medium of 90% Fetal Bovine Serum (FBS) and 10% Dimethyl Sulfoxide (DMSO) for cryopreservation.

### Preparation of CSF Samples

CSF samples were centrifuged at 400xg for 15 minutes at 4^0^C. After centrifugation, CSF fluids were put into -80^0^C for further use and pellets were used to determine CD20^+^ NK cells.

### Culturing of K562 Cells

K562 Human Myeloid Leukemia Cell Line (ATCC, CCL-243) was cultured in DMEM medium containing 10% FBS and 1% Penicillin-Streptomycin (Gibco, USA) at 37^0^C 5% CO_2_ incubator for three days. Cells were harvested upon reaching ∼85% confluence and 5 × 10^6^ cells were used for downstream applications while remaining cells were cryopreserved in liquid nitrogen for further use.

### Immunophenotyping of CD56^+^CD20^+^ NK Cells

Presence of CD56^+^CD20^+^ cells were determined by flow cytometric immunophenotyping. PBMC, CSF and homogenized tissue samples were placed into three tubes. Cells in the first tube were used as unstained negative control. Second tube was stained with anti-human CD3, CD56, CD14, CD19 and Zombie NIR FVD whereas the third tube was stained with the full antibody panel. **Table S17** shows the brands, clones and conjugated fluorochromes of antibodies used in the experiment. Samples were incubated with antibodies for 20 minutes at dark and washed with 2ml of staining buffer at 500g for 5 minutes. Supernatant was discarded and cells were resuspended with 1ml of staining buffer and analyzed under Attune NxT Focusing Cytometer (Thermo Scientific, USA). 1×10^6^ cells for PBMCs and all of the cells in CSF were analyzed within the Lymphocyte gate in FSC X SSC plot. FlowJo v.10.10.0 (BD Biosciences USA) was used to analyse the results.

### Functional Analysis of CD56^+^CD20^+^ Cells

Ten cryopreserved PBMCs were thawed at 37^0^C water bath, taken into complete medium (RPMI 1640 ^+^ 10% FBS ^+^ 1% Penicillin-Streptomycin, GIBCO, USA) and rested overnight 37^0^C 5% CO_2_ incubator. CD56^+^ NK Cells were isolated from PBMCs by using MojoSort Human CD56^+^ NK Isolation Kit (Biolegend, USA) according to manufacturer’s instructions. Isolated CD56^+^ cells were seeded into 96 well round bottom plates at a density of 1 × 10^5^ cells/100 ul and stimulated with Cell Activation Cocktail (PMA/Ionomycin without Brefeldin A, Biolegend, USA), IL-2 (10ng/ml) and IL-15 (10ng/ml) (Biolegend, USA) and 1 × 10^5^ K562 Cells/100ul with IL-2/IL15 for 1 hour at 37^0^C 5% CO_2_ incubator, respectively. After 1 hour, Brefeldin A (Biolegend, USA) was added into the wells and an additional stimulation was carried out for 5 hours. Two sets of antibody panels were used to determine the functions of CD20^+^ NK Cells which are given in **Table S18**. Results were analysed by FlowJo v.10.10.0 (BD Biosciences USA).

### K562 Killing Assay

1×10^4^ CD20^+^ and CD20^−^ NK cells from PBMCs of healthy controls were sorted with FACS Aria III Cell Sorter (BD Biosciences, USA) and cell purity and viabilty were determined afterwards. The experiment was continued if <95% purity and viability were achieved. K562 cells were labeled with CFSE (Biolegend, USA) according to the manufacturer’s instructions. Sorted NK cells and labelled K562 cells were co-cultered in a 96-well round bottom plate at a 2.5:1 effector to target ratio in complete RPMI medium for six hours in a 37^0^C 5% CO_2_ incubator with the presence of recombinant human IL-2 (10 ng/ml) and IL-15 (10 ng/ml) (Biolegend, USA). After incubation, samples were centrifuged once at 500xg for 5 minutes. Supernatants were collected and later used for multiplex cytokine analysis. Cells were stained with Annexin-V PE Apoptosis Detection Kit with 7-AAD (Biolegend, USA) according to manufacturer’s instructions.

CFSE stained K562 cells were gated, and apoptosis was determined by measuring only Annexin-V for early apoptosis, Annexin-V^+^7AAD^+^ for late apoptosis and 7AAD^+^ for non-apoptotic cell death. Materials used in the study is given in **Table S19**. Results were analysed by FlowJo v.10.10.0 (BD Biosciences USA).

### Measurement of inflammatory molecules secreted by NK cells

Supernatants collected during the K562 killing assay were used to determine the concentration of TNF-α, IFN-γ, granzyme A, granzyme B, perforin, granulysin, sFas and sFasL molecules secreted from CD20^+^ and CD20^−^ NK Cells respectively by using 13plex LegendPlex Human CD8/NK Kit (Biolegend, USA) according to manufacturer’s instructions. Briefly, standards and samples were combined with APC conjugated premixed beads coated with capture antibodies in a 96 well round bottom plate and incubated at RT for 2 hours. After incubation, plate was centrifuged with wash buffer and beads were resuspended with detection antibodies for 1 hour. Streptavidin-PE was added onto the wells and plate was incubated for another 30 minutes. Finally, plate was centrifuged and resuspended with 200ul wash buffer. Resuspended beads were transferred into 12×75 mm 5ml tubes and additonal 300ul wash buffer was added onto them. Cytokine standards and samples were run under Attune NxT Acoustic Cytometer (Thermo Fisher, USA). Beads were gated with FSC vs SSC plot whereas APC vs PE plot was used to classify beads and record MFI values for each cytokine, respectively. Collected data was analyzed with LegendPlex Analysis Software v.8.0 (Biolegend, USA).

### NK cell proliferation assay

PBMCs from 8 healthy control subjects were included in the NK proliferation studies. 1×10^7^ PBMCs were stained with CFSE Cell Division Tracker Kit (Biolegend, USA) according to manufacturer’s instructions. After staining, 1×10^6^ cells were seeded into 96 well U-Bottom plates as Day-2, Day-4, and Day-7 with unstimulated and stimulated wells. 10 ng/ml IL-2 and 10 ng/ml IL-15 (Biolegend, USA) were added to stimulated wells and plates were placed in a 37^0^C 5% CO_2_ incubator. Samples were stained after 2nd, 4th, and 7th days with Zombie NIR Fixable Viability Dye, mouse anti-human CD20 PE (clone 2H7), CD19 APC (clone HIB19), CD56 PE-Cy5 (clone MEM-188), CD3 APC-Cy7 (clone SK7), CD14 APC-Cy7 (clone HCD14). All antibodies and FVDs were purchased from Biolegend (Biolegend, USA). Acquisition was done with Cytoflex SRT (Beckman-Coulter, USA) and proliferation modelling analysis were done by FlowJo v.10.10.0 (BD Biosciences, USA).

### Single-cell RNAseq Data Analysis

All datasets are analyzed using the Scanpy pipeline.

### Twin MS dataset

The study by Ingelfinger et. al. contains peripheral blood samples from twins concordant with MS (*11*). Study is composed of two sorting schema, CD3^+^CD4^+^ T helper cells and CD3^−^CD11c^+^ myeloid cells. The same quality control filters with the original article were applied. Scrublet is used for doublet detection (*23*). The dataset is normalized to 10000 and then log(x+1) transformation is applied. Principal component analysis has been computed after dataset is scaled to zero mean and unit variance using Scanpy’s PCA function with arpack solver. Batch correction is performed with harmony algorithm on the principal components and the subsequent neighborhood graph is computed on the first 50 harmony-corrected principal components (*24*). Clustering is performed with Leiden algorithm by setting the resolution to 0.3 to catch main NK subtypes (*25*).

### COVID dataset

The dataset of the COMBAT consortium was downloaded as the processed AnnData object from Zenodo (*12*). NK cells are selected based on the original paper’s annotations. Then DGE analysis is conducted using Wilcoxon rank-sum statistics with Benjamini-Hochberg correction. Calculations are carried with “sc.tl.rank_genes_groups” function of the Scanpy. Since this data is created with the CITE-seq protocol, both antibody-derived tags (ADT) and RNA were available. Differential gene expression analysis is conducted for both RNA and ADT assays. CD20 expression of cells is determined using RNA assay and the cells were compared using this annotation for the ADT assay, as well. The differential expression analysis of both RNA and ADT assays are calculated using Wilcoxon rank-sum test.

### Melanoma dataset

The raw count matrix of the data by de Andrade et al. has been accessed from GEO with the accession number GSE139249 (*14*). During the preprocessing of the dataset, cells were removed if contain less than 200 genes and genes have been removed if expressed less than 3 cells. Doublets are identified and removed using Scrublet (*23*). Data is normalized to 1000 and log(x+1) is transformed. PCA is calculated on the data layer that is scaled to zero mean and unit variance. Batches are integrated using the Harmony algorithm. The neighborhood graph is computed by Scanpy using the default UMAP kernel on harmony-corrected principal component space. Contaminating cells have been removed in initial clustering through both manual inspection of clusters and automated cell type prediction using CellTypist’s “*Immune_High_All.pkl*” model (*15*). Additionally, the majority voting classifier is enabled to increase classification accuracy.

### Cross-tissue immune cell dataset

Data by Conde et. al. has been downloaded from the tissue immune cell atlas website (*15*). Preprocessed AnnData object is used for the downstream tasks. Since the additional training data for CellTypist was curated from the literature and the count matrix wasn’t available, we took another approach to remove B cells that may potentially contaminate NK cells. Gene set score is calculated using a set of B cell markers using “sc.tl.score_genes” function. The cutoff has been set to the zero and all cells with a B cell gene signature has been removed. After that, CD20^+^ vs CD20^−^ NK cells are compared using “sc.tl.rank_genes_groups” function as a validation. No B cell genes were upregulated among the CD20^+^ NK cell population.

### WGCNA Analysis

Weighted gene coexpression analysis (WGCNA) analysis was conducted on highly variable genes detected by Scanpy’s “*highly_variable_genes*” function. A signed hybrid network is conducted using the pyWGCNA package (*26*). Modules have been visualized with igraph network analysis software (*27*).

### Gene Set Enrichment Analysis

GSEA with hypergeometric enrichment test is performed with EnrichR using upregulated genes from differential gene expression analysis (*28*). “BioCarta_2016”, “GO_Biological_Processes_2023”, and “GO_Cellular_Components_2023” libraries were used for testing. Plots are generated using Seaborn and Matplotlib.

### Statistical Analysis

CD56^+^CD20^+^ cell population percentages were compared between HC, MS, IDD, OIND and NINS groups, functional, K562 cytotoxicity and cytotoxic molecule analysis between CD56^+^CD20^+^ and CD56^+^CD20^−^ cells as well as AIH PBMCs, BM, Tumor, and healthy tissue immunophenotyping analysis were done by Mann-Whitney U Rank-Sum test. Analysis in blood and CSF were analyzed by Spearman correlation. GraphPad v.8 was used as statistical analysis software (GraphPad Software, USA).

## Acknowledgements

We would like to thank to the patients and healthy volunteers who participated in this research project. We are also thankful to Prof. Dr. Caner Süsal and Dr. Miriam Fichtner for reviewing the first draft of our manuscript and their invaluable comments. The authors gratefully acknowledge the use of the services and facilities of the Koç University Research Center for Translational Medicine (KUTTAM), funded by the Presidency of Turkey, Presidency of Strategy and Budget. During the preparation of this work the authors used ChatGPT 3.5 for English language editing of this article. After using this tool, the authors reviewed and edited the content as needed and takes full responsibility for the content of the publication.

## Author contributions

Conceptualization: AV

Design of experiments: AV, ÖA

Investigation: ÖA, ET, NA, ABK

Formal analysis: AV, ÖA, ET

Resources: NA, ABK, TD, GG, MY, BU, MÜ, IB, KS, GE, IM, MK, BY, SE, ÇA, MZ, AS

Funding acquisition: AV, ÖA

Visualization: AV, ÖA, ET Supervision: AV, EM

Writing – original draft: AV, ÖA, ET

Writing – review & editing: GE, SV, MZ, EM

## Data and materials availability

All data are available in the main text or the supplementary materials.

