## Supplementary Figures for "CD20^+^ natural killer cells are polyfunctional, memory-like cells that are enriched in inflammatory disorders"

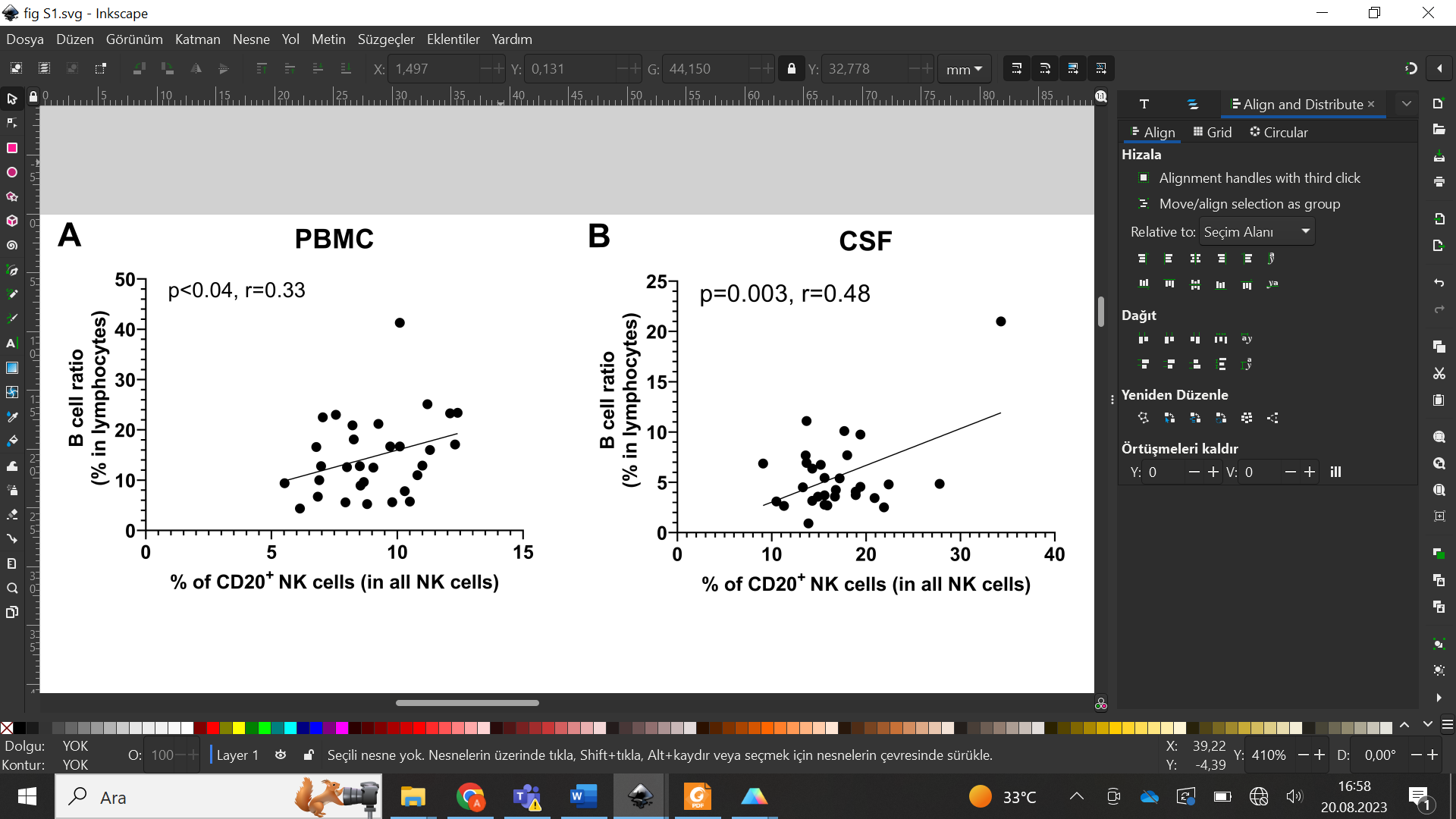

**Fig. S1.** The ratio of B cells is correlated with that of CD56^+^CD20^+^ cells in both peripheral blood and CSF of pwMS. P and r values are calculated by Pearson’s correlation test.

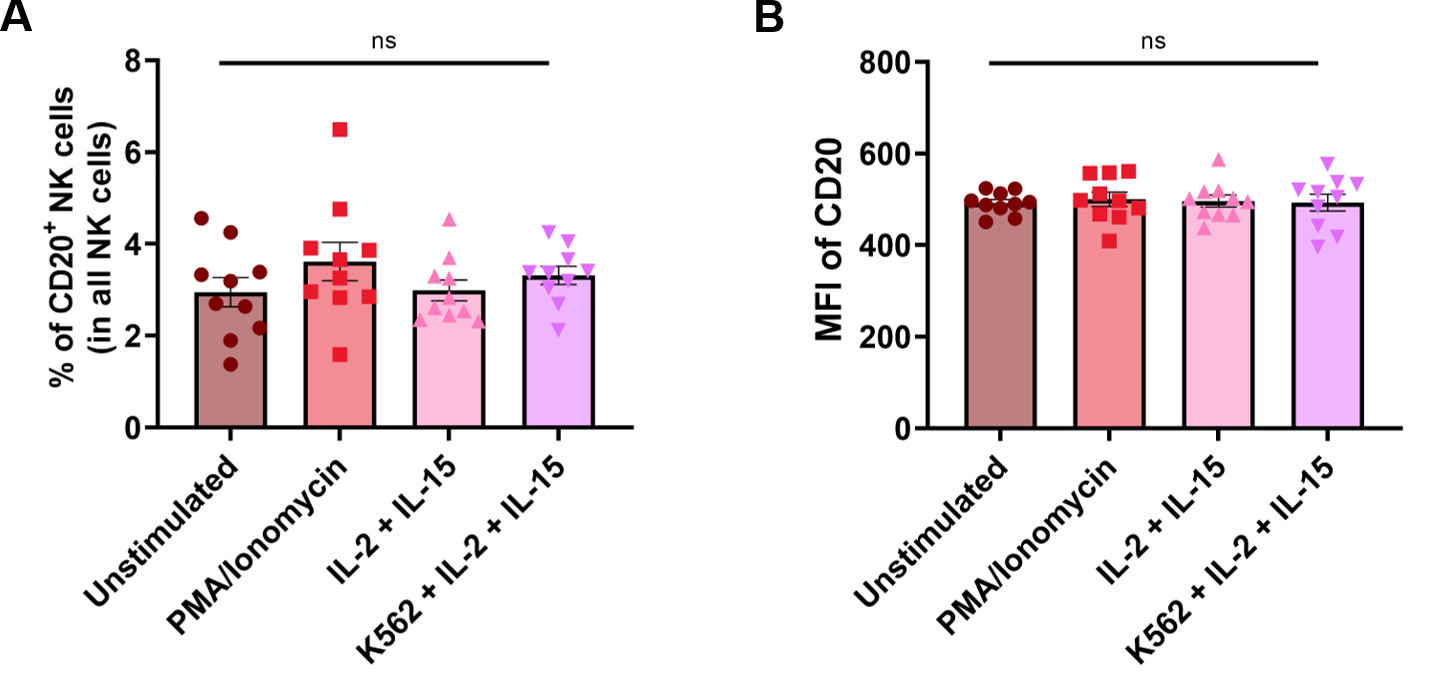

**Fig. S2.** The percentage and MFI of CD20^+^ NK cells are not changed after 6h of stimulation with PMA/I, IL2/IL15 or K562 plus IL2/IL15. (A) shows the percentage of CD56^+^CD20^+^ cells among CD56^+^ NK cells. (B) represents the expression level of CD20 on CD20^+^CD56^+^ cells after the indicated stimulation conditions.

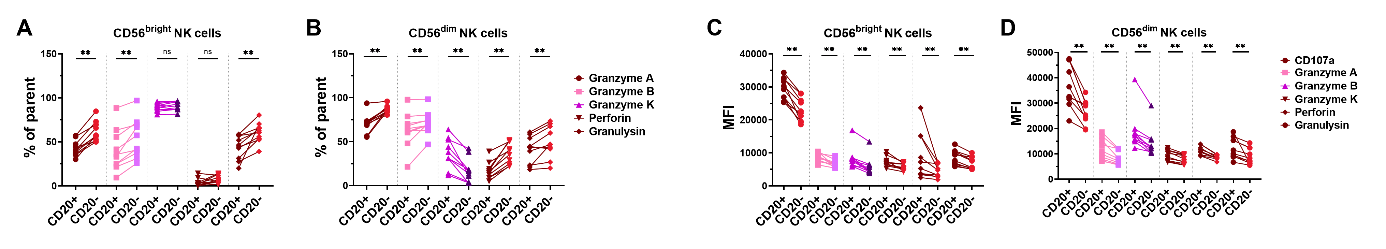

**Fig. S3.** Expression of the cytolytic molecules by CD20^+^ NK cells among the CD56^bright^ and CD56^dim^ NK cells (A and B) Percentage of cells. (C and D) MFI of cytotoxic molecules.

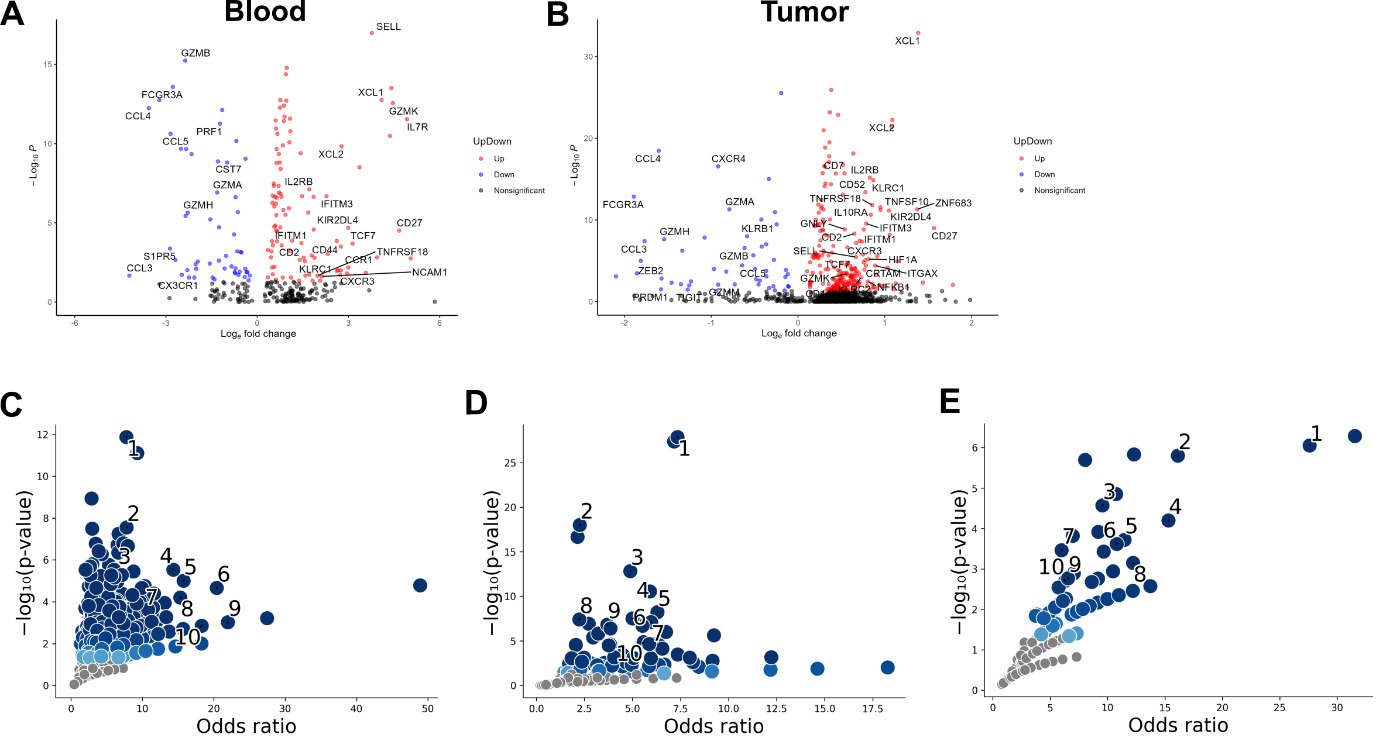
**Fig. S4.** Differential gene expression analysis of *MS4A1* expressing NK cells versus non-expressing NK cells in the blood (A) and tumor tissue (B) of melanoma patients. The graphs showing the p-value and odds ratio of the results of GO annotation analysis that are shown in figure 6 J and K (C and D, respectively), and gene set enrichment analysis shown in figure 6L (E).
